# Individual uniqueness of connectivity gradients is driven by the complexity of the embedded networks and their dispersion

**DOI:** 10.1101/2024.12.01.626248

**Authors:** Yvonne Serhan, Shaymaa Darawshy, Wei Wei, Daniel S. Margulies, Karl-Heinz Nenning, Smadar Ovadia-Caro

**Affiliations:** Department of Cognitive Sciences, School of Psychological Sciences, Faculty of Social Sciences, University of Haifa, Haifa, Israel; The Integrated Brain and Behavior Research Center (IBBRC), University of Haifa, Haifa, Israel; Centre National de la Recherche Scientifique and Université de Paris, INCC UMR 8002, Paris, France; Nathan S. Kline Institute for Psychiatric Research, Orangeburg, NY, United States

**Keywords:** resting-state fMRI, connectome fingerprinting, identity analysis, test-retest similarity, Dimensionality reduction

## Abstract

Connectivity gradients are widely used to characterize meaningful principles of functional brain organization in health and disease. However, the degree of individual uniqueness and shared common principles is not yet fully understood. Here, we leveraged the Hangzhou test-retest dataset, comprising repeated resting-state fMRI scans over the span of one month, to investigate the balance between individual variation and shared patterns of brain organization. We quantified the short- and long-term stability for the first three connectivity gradients and used connectome fingerprinting to establish the associated individual identification rate. We found that all three connectivity gradients are highly correlated over both short and long time intervals, demonstrating connectome fingerprinting utility. Individual uniqueness was dictated by the complexity of the networks such that heteromodal networks had higher connectome fingerprinting rates than unimodal networks. Importantly, the dispersion of the gradient coefficients associated with canonical functional networks was correlated with identification rates, irrespective of the position along the gradients. Beyond individual uniqueness, between subject similarity was high along the first connectivity gradient, which captures the dissociation between unimodal and heteromodal cortices, and the second connectivity gradient, which differentiates sensory cortices. Our results support the usage of connectivity gradients for the purposes of both group comparisons and prediction of individual behaviours. Our work adds to existing knowledge on the shared versus unique organizational principles and offers insights into the importance of network dispersion to the individual uniqueness it carries.

## Introduction

Studying individual differences in brain-behavior associations based on features of brain organization relies on the premise that some organizational features are unique to individuals, stable over time and can be used as a proxy for individual behaviors, individual symptomatology and individual recovery (Damoiseaux et al., 2006; Finn et al., 2015; Fornito & Bullmore, 2015; Gillebert & Mantini, 2013; Ramduny & Kelly, 2024; Shehzad et al., 2009). Understanding the intricate relationship between what is shared and what is unique to each individual constitutes a major challenge in neuroscience with both theoretical and practical implications.

The mapping of macroscale organization using intrinsic activity fluctuations measured with resting-state functional MRI (rs-fMRI) (Biswal et al., 1995) is offering a non-invasive means for representing both features of segregation into distinct functional modules as well as their integration by hyperconnected networks (Bertolero et al., 2018; Bullmore & Sporns, 2009; Cohen & D’Esposito, 2016; van den Heuvel & Sporns, 2011). Whereas a-priori parcellations are commonly used to overcome the high-dimensionality of rs-fMRI connectivity data (Gordon et al., 2016; Smith et al., 2009), more recently, data-driven, dimensionality reduction techniques are applied to represent a low-dimensional connectivity space, or connectivity axes, also termed *connectivity gradients* (Haak et al., 2018; Langs et al., 2014; Margulies et al., 2016).

The main strength of the approach is the continuity reflected in this representation along with the fundamental organizing principles it emphasizes. Brain regions are ranked along the axis of each connectivity gradient based on similarities in their whole-brain connectivity patterns (Huntenburg et al., 2018; Langs et al., 2014, 2016; Margulies et al., 2016), and different functional modules are therefore spread along each of the gradients in different locations yet in a clustered manner corresponding with well-known canonical functional networks (Hong et al., 2020; Huntenburg et al., 2018; Margulies et al., 2016). The fact that nodes are located along a continuum enables the adequate representation of both elements of segregation and integration. Whereas clustering of values reflects segregation and enhanced similarity of connectivity, integration is reflected in enhanced dispersion of values within a module, or an overlap of values between different modules. These changes in values correspond with ‘contraction’ or ‘expansion’ of the gradient as referred to in previous work (Knodt et al., 2023; Nenning et al., 2020; Ottoy et al., 2024). It is therefore that the *dispersion* of values is a mirror of both the modular topology and the connectedness of the nodes, or the network they are affiliated with.

From the representational perspective, the first connectivity gradient which is the one commonly used is reflecting a hierarchy such that primary sensory and unimodal networks are clustered at one end, whereas high-order, heteromodal networks and the default-mode network are at the other end (Margulies et al., 2016, 2022). This eminent orderly organizing principle (M. M. Mesulam, 1998) emerging in a data-driven manner further supports the biological validity of the approach and provided the initial empirical support for its wide-usage. Beyond the first connectivity gradient, the second and third connectivity gradients usually emphasize different networks on their extremes (Bayrak et al., 2019; Margulies et al., 2022). Whereas the second connectivity gradient distinguishes the visual network from most other networks, the third connectivity gradient is demonstrating a different arrangement of networks such that areas of the frontoparietal and dorsal attention networks are situated at one end and areas of the default-mode network are situated on the other end. Different gradients therefore emphasize networks of different complexities on their extremes as well as different degrees of segregation or integration of these networks. Whereas along the first gradient, these two features correspond with one another as a continuum from unimodal to heteromodal, along the second gradient, a more subtle hierarchy within the visual network is captured. Gradients therefore contain both information on networks’ complexity, or their hierarchical role (unimodal/heteromodal) in addition to their range of values, or dispersion, i.e., the degree to which they integrate via shared connectivity patterns.

Considering the advantages stated above, multiple studies in recent years have utilized connectivity gradients for both group comparisons and prediction of individual behaviour (Bayrak et al., 2019; Bernhardt et al., 2022; Bethlehem et al., 2020; Brown et al., 2022; Dong et al., 2023; Girn et al., 2022; Haak et al., 2018; Hong et al., 2019; Huntenburg et al., 2018; Knodt et al., 2023; Meng et al., 2021). However, the degree of similarity and the exploration of shared versus unique features captured along different gradients have not been directly investigated beyond the first connectivity gradient (Hong et al., 2020; Knodt et al., 2023). Since some studies are using multiple gradients and some even provide measures based on the combination of different gradients (Bethlehem et al., 2020), information on the shared and unique features of different gradients is essential to make informed analysis decisions fitted to the question at hand.

Here, we aimed at assessing the stability of connectivity gradients and their individual uniqueness using a subset of a freely-available test-retest data collected in Hangzhou University (B. Chen et al., 2015). Since individual test-retest reliability has been shown to decrease as intervals between the scans increase in multiple studies (Birn et al., 2013; Fiecas et al., 2013; Noble et al., 2017; O’Connor et al., 2017; Pannunzi et al., 2017; Shehzad et al., 2009), we chose to focus on both short-term interval (two scans separated by three days) and long-term interval (two scans separated by one month). Given the wealth of data supporting the functional significance of connectivity gradients in both group studies and as predictors of individual behaviors (Bayrak et al., 2019; Bernhardt et al., 2022; Bethlehem et al., 2020; Brown et al., 2022; Dong et al., 2023; Girn et al., 2022; Haak et al., 2018; Hong et al., 2019; Huntenburg et al., 2018; Knodt et al., 2023; Meng et al., 2021; Xia et al., 2022), as well as previous work addressing reliability of connectivity gradients (Hong et al., 2020; Knodt et al., 2023), we hypothesize that connectivity gradients will demonstrate high spatial similarity between different subjects, particularly for the first connectivity gradient. We additionally expect that connectivity gradients will be individually unique such that they will show high similarity within subjects, sufficient for effective connectome fingerprinting (Ramduny & Kelly, 2024; Finn et al., 2015). As for the factors influencing the effective fingerprinting, we hypothesize that the complexity of a network along the hierarchy and its dispersion will drive individual uniqueness. This hypothesis is supported by both empirical work showing high-order networks dominate individual uniqueness in parcellation-based studies (Finn et al., 2015; Blautzik et al., 2013; Marchitelli et al., 2017; Mejia et al., 2018; Noble et al., 2017, 2019; O’Connor et al., 2017; Somandepalli et al., 2015; Wisner et al., 2013; Zuo et al., 2014), as well as more theoretical accounts on the link between enhanced range and complex behaviors, where behavioral diversity is most likely to be expressed (M. Mesulam, 2011; M. M. Mesulam, 1998; M.-M. Mesulam, 1990).

## Methods

### Subjects’ information

The current study is using a subset of the Hangzhou online available test-retest data (B. Chen et al., 2015). Thirty participants (age range 20-30 years, 15 females, mean age = 24, SD = 2.41) were recruited and repeatedly scanned over one month. Here, we used three consecutive rs-fMRI sessions from this dataset. The first two sessions were collected three days apart and the third session was collected 30 days after the first session (referred to here as days 1, 3, and 30). We used these three time points to investigate similarity over short and longer time intervals. Data collection was approved by the ethics committee of the Center for Cognition and Brain Disorders (CCBD) at the Hangzhou Normal University. Written informed consent was obtained from all participants.

### MRI data acquisition

MRI imaging was performed using a GE MR750 3.0 Tesla scanner (GE Medical systems, Waukesha, WI) at CCBD at Hangzhou Normal University. Anatomical and functional MRI data were collected at each time point. A T1-weighted Fast Spoiled Gradient echo (FSPGR: TR = 8.06 ms, TE = 3.1 ms. TI = 450 ms, flip angle = 8 degrees, field of view = 256 * 256, voxel size = 1 * 1 * 1 mm, 176 sagittal slices) was performed to acquire a high-resolution anatomical image. A 10-min T2-weighted echo-planar imaging (EPI: TR = 2000 ms, TE = 30ms, flip angle = 90 degrees, field of view = 220 * 220 mm, matrix = 64 * 64, Voxel size = 3.4 * 3.4 * 3.4, 43 slices) sequence was performed to obtain functional images. The subjects had their eyes open, and a screen presented a black fixation point ‘+’ in its centre. The participants were instructed to relax and remain still with their eyes open, not to fall asleep, and not to think about anything in particular (Chen et al., 2015).

### MRI data preprocessing

Imaging data were preprocessed using fMRIPrep 20.2.3 (Esteban et al., 2019; Esteban, Blair, et al. (2018)). Anatomical preprocessing included: intensity non-uniformity (INU) correction, skull-stripping of the T1-weighted (T1w) reference image, and segmentation of cerebrospinal fluid (CSF), white matter (WM), and gray matter (GM). Reconstruction of brain surfaces was applied using recon-all from FreeSurfer 6.0.1. Spatial normalization to the MNI standard space (MNI152NLin2009cAsym) was performed using nonlinear registration with antsRegistration, using both brain-extracted anatomical and functional reference images. Functional preprocessing included the following steps for each of the three sessions per subject: motion correction, slice scan time correction, extraction of signals from the white matter and CSF masks, and extraction of a reference volume for functional to T1 registration by computing median over motion corrected file. The denoising strategy was performed using load confound python package (https://github.com/SIMEXP/load_confounds) and included a minimal denoising strategy: regression of full motion parameters (6 parameters), white matter, CSF, and additional high pass filter (0.01Hz). The final file was transformed to a fsaverage5 surface template using mri_vol2surf (FreeSurfer). Correlation matrix was then calculated for each MNI normalized dataset, resulting in 20,484 * 20,484 entries. This correlation matrix was fed to the dimensionality reduction algorithm.

### Connectivity gradients analysis

Connectivity gradients represent axes of variance in the data by using techniques such as decomposition or embedding, which reduce the original dimensions while preserving most of the variability of the original data (Huntenburg et al., 2018). Each component therefore contains loading coefficient values assigned to each vertex, indicating the similarity of global connectivity patterns among vertices. Specifically here, for each scan, time series files were extracted for each vertex and correlation matrix was computed resulting in a 20,484*20,484 correlation matrix. Individual correlation matrices were fed to the dimensionality reduction algorithm, Principal Component Analysis (PCA) (Hong et al., 2020) resulting in 5 components/gradients for each subject and each scan. Gradient vectors were then aligned to a common template (Margulies et al., 2016) using the generalized Procrustes alignment approach (Vos de Wael et al., 2020). This choice of using only the first five components was based on previous work demonstrating they account for most of the variance in the original data, and work demonstrating that adding more components does not significantly enhance the explained variance for subsequent analysis (Bethlehem et al., 2020). Due to the predominant focus on the first gradient in previous research (Bernhardt et al., 2022) and the inherent interpretability challenges associated with multiple gradients, we have limited our analysis to the first three gradients. This decision is consistent with our previous work, which demonstrated that the first and third connectivity gradients serve as a biologically valid model (Bayrak et al., 2019), and with other studies that have identified behaviorally meaningful effects using these gradients (Hong et al., 2019) (see supplementary Fig S1 for explained variance as a function of gradient number).

### Similarity and individual identification analysis

To quantify how similar subjects are to themselves, Identification analysis, also known as connectome fingerprinting analysis was conducted similarly to previous work (Finn et al., 2015; Ramduny & Kelly, 2024). First, to quantify stability over repeated tests, spatial similarity was calculated for each gradient using Pearson’s correlation coefficient. Correlation was computed over vector pairs (day 1 - day 3 and day 1 - day 30) for all subjects yielding spatial similarity matrices for each session pairs (short and long-term) and for each connectivity gradient. Fingerprint analysis assigns an ID based on the highest matched correlation value (a ‘winner take all approach’). Identification rates were therefore calculated as the proportion of correct classification of the same subjects as themselves over two sessions. To assess the statistical significance of identification rates, a non-parametric permutation test was then performed. In each iteration, subjects ID was permuted such that a ‘correct’ classification was assigned to a different subject to create a null distribution of ID accuracies. 1000 iterations were employed as previously done (Finn et al., 2015).

To assess stability over repeated measurements within and between subjects, correlation was averaged for within-subjects values (along the diagonal) and between subjects values. A ratio of within/between subjects was calculated to reflect stability within individuals while taking into account correlation between subjects. We examined how this ratio varied across different gradients and whether it changed between sessions. A repeated-measures ANOVA was employed to assess the effect of the different gradients and different time on this measure. The model included a subject-specific random effect to account for within-subject variability, with subjects as the error term. The full model examined the main effects of connectivity gradient and time, as well as their interaction. Additionally, post-hoc pairwise comparisons between the gradient levels were performed using the Tukey method to adjust for multiple comparisons. All statistical analyses were performed using R, and statistical significance was determined at an alpha level of 0.05.

### A-posteriori parcellation to networks

To address the relative contribution of different functional networks to the observed accuracy, a-posteriori parcellation (Thomas Yeo et al., 2011) to 6 canonical cortical networks was employed. The limbic network was omitted from our analysis due to low signal-to-noise ratio in these areas, which may inaccurately reflect connectivity among its regions through its wide range of values (Thomas Yeo et al., 2011). Loading values along each of the connectivity gradients were extracted for each network and ID analysis was then performed based on ICC obtained from each network using only the relevant vertices in each parcel. To explore additional unimodal networks beyond sensorimotor and visual domains represented in Yeo’s parcellation, the Cole-Anticevic Brain-wide Network Partition was additionally used in the same manner (Ji et al., 2018). This process resulted in the subdivision of the six networks into subsystems, yielding a total of 12 networks. This expansion encompasses the distinction between primary and secondary visual regions, as well as the inclusion of the auditory network. Ventral Multimodal (VMM) and Orbito-affective (ORA) parcels were excluded similarly to the approach we applied using Yeo’s parcellation due to low signal-to-noise ratio and small network size for these two networks (Sanders et al., 2022). A total of 10 subnetworks was therefore used for further analyses.

### Network’s complexity and dispersion

The networks were categorized into two groups: heteromodal and unimodal networks across both parcellations. To assess whether there were differences in accuracy rates between these groups, a Wilcoxon signed-rank test was performed in each parcellation separately. To determine whether changes in variance between sessions and gradients influenced the accuracy of the ID analysis, the variance in loading values was calculated for each network. Variance, in this context, serves as a measure of the dispersion of gradient loading values within a specific network. Higher variance indicates a broader range, i.e., larger distribution of loading values and more integration along the continuum. Conversely, lower variance suggests a more uniform distribution, or a segregated connectivity pattern. A partial correlation analysis was performed to assess the relationship between variance and ID accuracy while controlling for the number of vertices in each network.

## Results

### Gradient maps represent connectivity space on three axes based on shared connectivity patterns

To represent connectivity data on a lower-dimensional space, the first three connectivity gradients were extracted for each dataset individually. Following the application of PCA, each vertex receives a unitless loading value. Along each one of the gradients, vertices that share similar loading values, share a similar global connectivity pattern. As a result, functional networks tend to cluster along specific gradients. The different gradients, however, implicate different networks on their extremes and therefore capture a different representation. Average group-maps over all subjects and all sessions (Fig.1a) was calculated and an a-posteriori parcellation to six canonical networks (Thomas Yeo et al., 2011) was employed to visualize representations of the different networks and their location along the three gradients. Along the first gradient, heteromodal regions, such as the default mode network (DMN), exhibit high loading values and are clustered at one end of the gradient. In contrast, unimodal regions, including the sensorimotor network (SMN) and the visual network (VN), are positioned at the opposite end, reflecting a transition from unimodal to heteromodal networks. This hierarchical organization is consistent with the gradient structure previously reported (Margulies et al., 2016; Nenning et al., 2020), also for studies using different decomposition approaches (Bayrak et al., 2019; Hong et al., 2020). Different distinction emerges along the second gradient, where the visual network (VN) is segregated from the sensorimotor network (SMN) and most other cortical regions. This differentiation highlights the dissociation of the visual system from most other functional networks. The third gradient captures a separation between different networks involved in higher-order cognitive functions. The frontoparietal network (FPN), dorsal attention network (DAN), and salience network (SN) are clustered together at one end, while the DMN is located at the opposite end of the gradient (Fig. 1b). The dispersion of values depicted in the density plots and the location of known networks changes in the different axes. The three connectivity gradients therefore capture different representations of functional networks and their respective segregation or integration along the continuum as reflected in the range of loading values. While this general layout is preserved in individual data demonstrating stability (Fig. 1c), some variation can still be observed over time thereby representing changes in the connectivity space within the same individual.

**Fig. 1.**
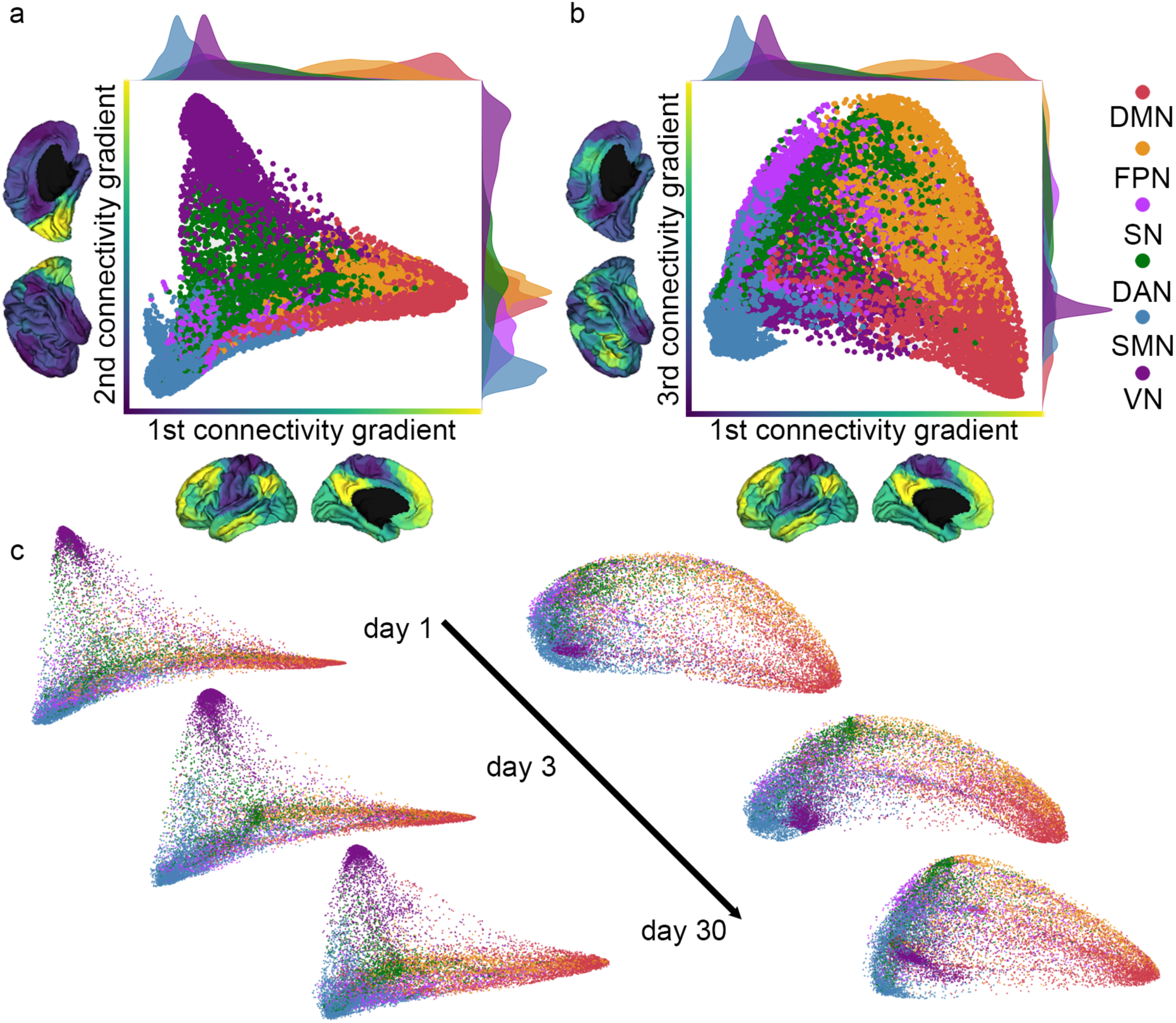
Connectivity gradients represent a continuum based on similarity of functional connectivity patterns. (a) Average loading values for all subjects for each vertex, color coded according to Yeo’s parcellation (Thomas Yeo et al., 2011). Along the first connectivity gradient (x-axis), a hierarchical gradient is captured such that default-mode network (DMN) is on the one end and sensorimotor network (SMN) and visual network (VN) are on the other end. Along the second connectivity gradient (y-axis) VN is distinguished from the SMN and most other networks. (b) Along the third connectivity gradient (y-axis), fronto-parietal network (FPN), dorsal attention network (DAN), and salience network (SN) are on the one end and DMN is on the other end. Different functional networks are represented in different locations along the different gradients and each gradient captures a different representation. Along the different gradients, networks can change their dispersion of coefficient values. (c) Example data from an individual subject scanned over time. Connectivity space is represented along the first and second connectivity gradients (left), and along the first and third connectivity gradients (right). The general structure captured in the connectivity space is retained showcasing stability yet slight changes are still evident over time.

### Connectivity gradients are stable within individuals over short and long-term periods

To quantify the stability of connectivity gradients over repeated measurements, Pearson’s Correlation Coefficient was used to quantify spatial similarity. Correlation values were calculated between session pairs at day 1-day 3 and day 1-day 30, both within and between subjects, resulting in a separate correlation matrix for each gradient. The diagonal of the matrix shows correlation values for individual subjects across the different sessions (Fig. 2). Correlation values along the diagonal were generally higher than correlation values obtained for different subjects. All three connectivity gradients demonstrated ‘high’ correlation values (mean r values > 0.7, Fig. 3a) (Mukaka, 2012) within individuals over both short and long time intervals. To examine if connectivity gradients can be effectively used for connectome fingerprinting, individual identification analysis (Finn et al., 2015) was performed. We found that for the first gradient accuracy rates were 83.34% for day 1 - day 3, and 80% for day 1 - day 30 cross prediction. For the second gradient, accuracy rates were 66.67% for day1 – day 3, and 40.34% for day 1 – day 30. For the third gradient, accuracy rates were 73.34% for day 1 – day 3, and 70% for day 1 – day 30. To assess the statistical significance of the accuracies, a nonparametric permutation test was performed. Among all iterations, the highest success rate observed was 6 out of 30, or roughly 20%, occurring in only 2 out of the 1,000 iterations. Hence, the p-value associated with the lowest accuracy we found is 0. Even though accuracy rates are slightly lower for longer time intervals (30 days apart) than for short time intervals (3 days apart), all three connectivity gradients were shown to be reliable predictors of individual organization, with the first connectivity gradient showing the highest accuracy rates in identification analysis.

**Fig. 2.**
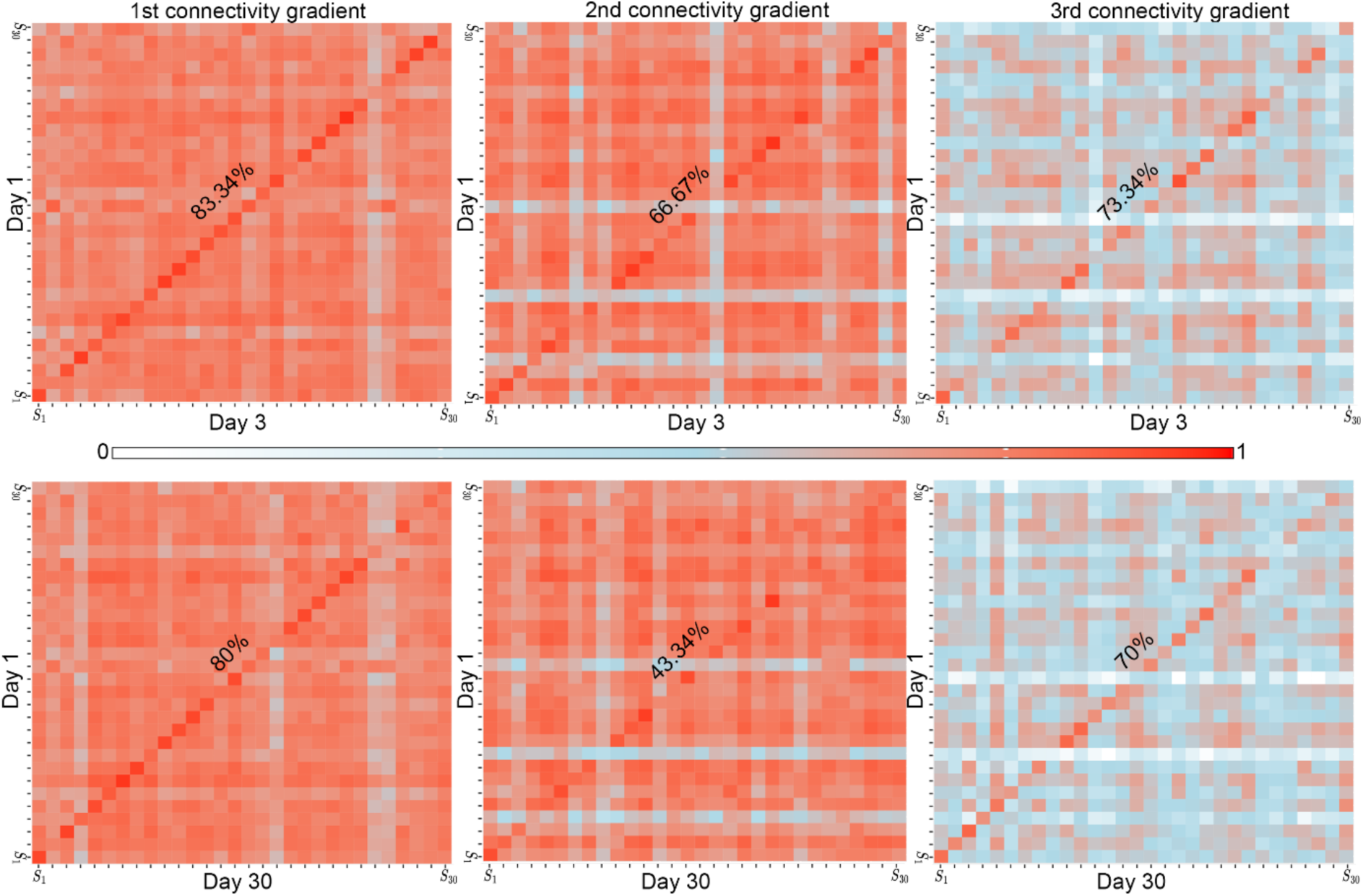
Connectivity gradients are stable within individuals over short and long-term intervals. (a) Pearson’s Correlation Coefficient values for connectivity gradients across subjects’ data pairs. Along the diagonal, all three gradients show ‘high’ correlation values (mean r values > 0.7), indicating high stability within subjects. In the identification analysis, the first gradient demonstrates the highest accuracy rates with 83.34% for Day 1 - Day 3, and 80% for Day 1 - Day 30. The second gradient showed slightly lower accuracies: 66.67% for Day 1 - Day 3, and 43.34% for Day 1 - Day 30. The third gradient shows accuracies of 73.34% for Day 1 - Day 3, and 70% for Day 1 - Day 30. Connectivity gradients are therefore individually unique in both short and long-term intervals for all three connectivity gradients.

**Fig. 3.**
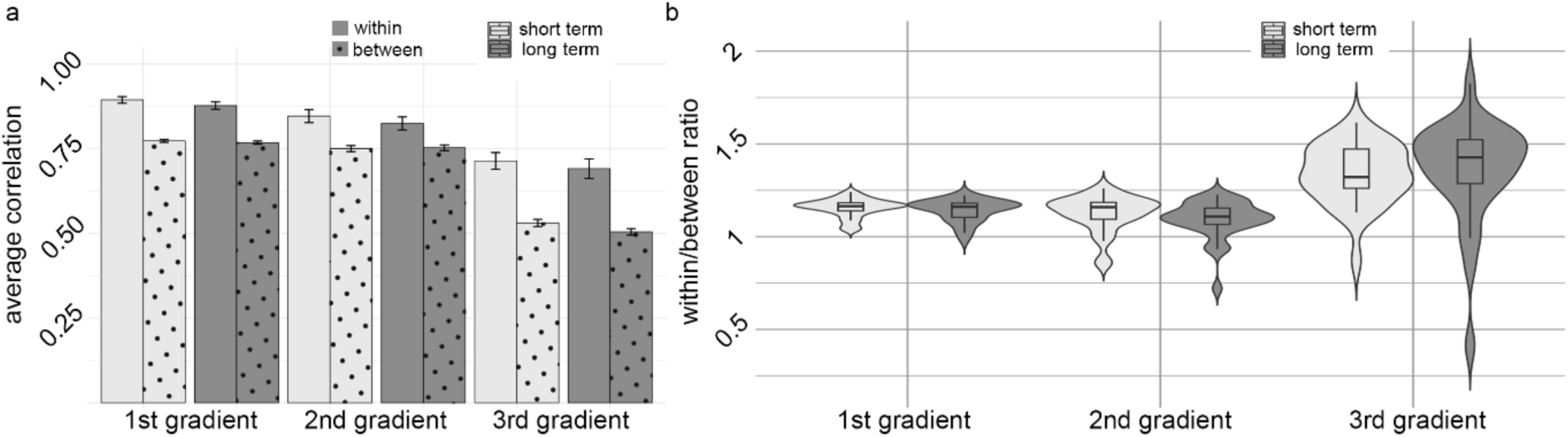
Within/between correlation values for the three connectivity gradients over short and long intervals. (a) Average Pearson’s correlation values for short-term (day 1 - day 3) and long-term (day 1 - 30 days) sessions across the first, second, and third gradients, within (plain) and between (dotted) subjects. Bars represent mean values with standard error. Between subjects’ correlation values are lower than within subjects’ correlation values, yet are high along the first and second connectivity gradients and moderate along the third connectivity gradient. Within subjects’ correlation values are high along all three gradients and time intervals. (b) Within-to-between ratio distributions for short and long intervals across gradients. Violin plots display the distribution, box plots show the median and interquartile range, and whiskers represent variability. A value above 1 indicates higher correlation values for within-subject correlations compared to between-subject correlations. All three gradients show a mean ratio above 1. Significant differences were found between the first and third gradient and between the second and the third gradient.

### The first and the second connectivity gradients are largely shared across different subjects

Beyond the high correlation values captured within the same subjects when re-scanned, considerable similarity exists between different subjects. Average correlation values between different subjects are in accordance with ‘high’ correlation for the first gradient and for the second gradient (mean r values > 0.7). Along the third gradient correlation drops to ‘moderate’ (Mukaka, 2012) (mean r values = 0.5) (Fig. 3a). To examine the relative contribution of within subject similarity taking between subjects’ similarity into account, a ratio of within/between correlation was calculated (Fig. 3b). A significant main effect for ‘gradient’ was found in repeated measures ANOVA (F(2,58)=52.11, p<.001), indicating significant differences across gradients. No significant main effect was observed for ‘time’ (F(1,29)=0.258, p=0.615), nor for interaction between ‘gradient’ and ‘time (F(2,58)=0.714, p=0.494). Post-hoc comparisons using the Tukey method revealed significant differences between Gradient 1 and Gradient 3 (t(115)=−5.879, p<.0001), and between Gradient 2 and Gradient 3 (t(115)=−7.274, p<.0001). No significant difference was found between Gradient 1 and Gradient 2 (t(115)=1.395, p=0.347).

### Higher complexity of functional networks is associated with higher accuracy rates in identification analysis

To investigate the contribution of specific resting-state networks (RSNs) to the high identification accuracy rates observed at the whole gradient level, a-posteriori parcellation (Thomas Yeo et al., 2011) was applied to the three connectivity gradients (Fig. 4a) and identification analysis was repeated using loading coefficient values in each pre-defined network. Accuracy rate were compared for heteromodal networks, including the dorsal attention network (DAN), frontoparietal network (FPN), default mode network (DMN), and salience network (SN), with unimodal networks, including the somatomotor network (SMN) and visual network (VN). Heteromodal networks demonstrated significantly higher accuracy rates along the first (t=4.88, p=0.0033) and third (t=7.64, p=8.87e−05) connectivity gradients. In contrast, along the second connectivity gradient, the visual network achieved moderate accuracies within the range of accuracies shown for other heteromodal networks and there is no significant difference between unimodal and heterometal networks (t-test: t=0.51, p=0.6332). To complement our analysis with an additional, more detailed parcellation the same analysis was repeated using Cole parcellation (Fig. 4b). Similar trends were observed, where heteromodal networks, such as the Cingulo-Opercular Network (CON), Posterior Multimodal Network (PMM), and Language Network (LAN), demonstrated higher accuracy rates as compared with unimodal networks along the first (t=9.514, p<0.001) and third (t=6.212, p<0.001) connectivity gradients, and not along the second connectivity gradient (t=−0.374, p=0.7141). Interestingly, along the second connectivity gradient, the Visual 2 (VIS2) network achieved relatively high accuracies (76.67% for Day 1 - Day 3, and 66.67% for Day 1 - Day 30), while lower-order visual areas, Visual 1 (VIS1), showed lower accuracies (43.34% for Day 1 - Day 3, and 56.67% for Day 1 - Day 30). This distinction between higher-order and lower-order visual regions accounts for the high accuracy rates obtained along the second gradient for the visual network. These findings replicate and extend previous work (Finn et al., 2015; Blautzik et al., 2013; Marchitelli et al., 2017; Mejia et al., 2018; Noble et al., 2017, 2019; O’Connor et al., 2017; Somandepalli et al., 2015; Wisner et al., 2013; Zuo et al., 2014), supporting the role of high-order heteromodal networks in individual connectome fingerprinting.

**Fig. 4.**
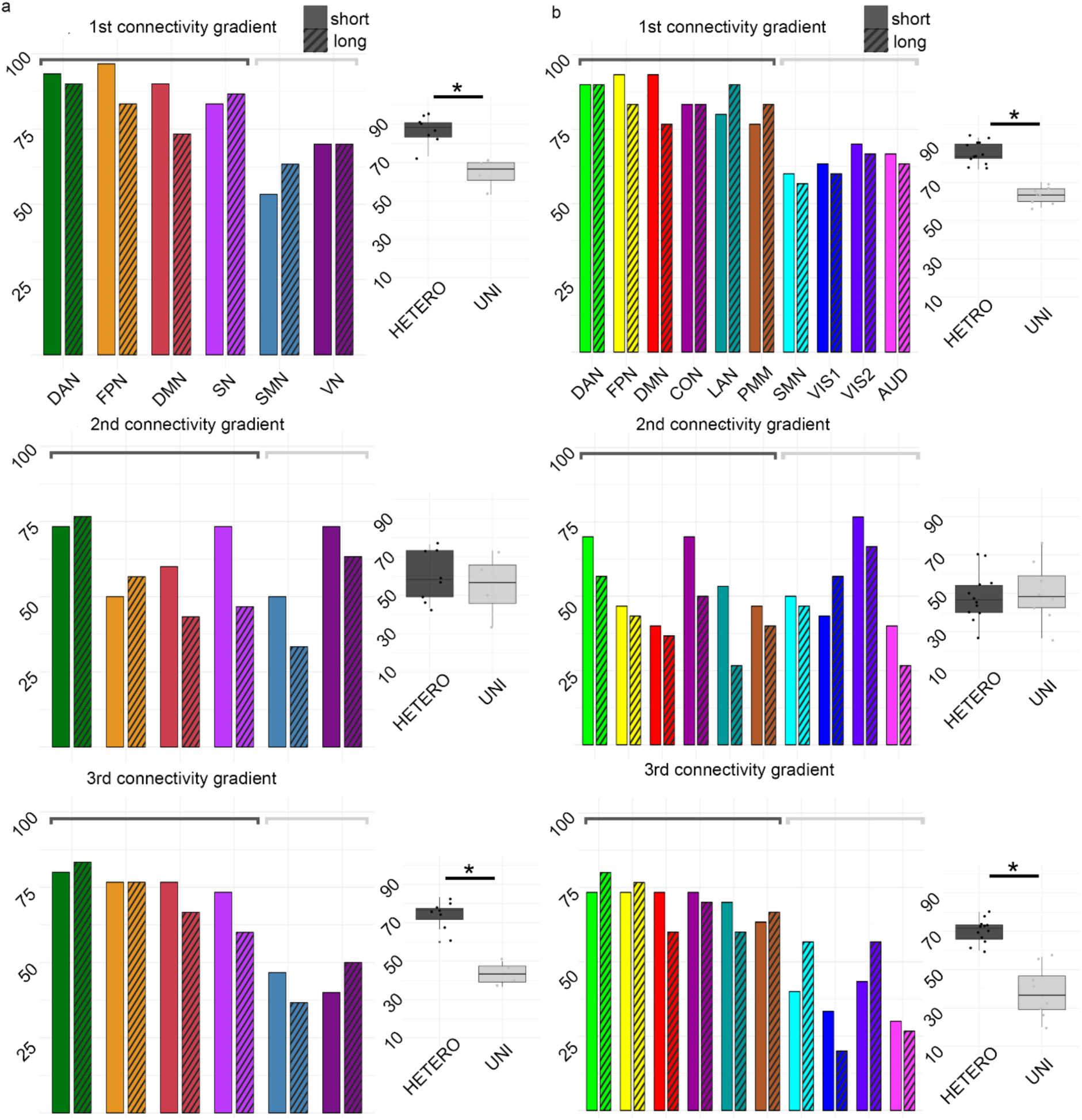
Networks of higher complexity show higher accuracies in individual identification analysis. Accuracy rates for short-term (dashed bars) and long-term (plain bars) sessions across different brain networks for the first three connectivity gradients (left). Box plots comparing accuracy rates for heteromodal (HETERO) and unimodal networks (UNI) (right). (a) Accuracy rates for Yeo parcellation, including the Dorsal Attention Network (DAN), Frontoparietal Network (FPN), Default-mode Network (DMN), Salience Network (SN), Somatomotor Network (SMN), and Visual Network (VN). (b) Accuracy rates for Cole parcellation, incorporating the Cingulo-Opercular Network (CON), Language Network (LAN), Posterior Multimodal (PMM), Visual 2 (VIS2), Visual 1 (VIS1), and Auditory Network (AUD). Along the first and third gradients, networks of higher complexity show consistently higher rates of individual fingerprinting. Interestingly, along the second connectivity gradient, the visual network achieves high accuracy, but there is no significant difference between unimodal and heteromodal networks. A closer look shows that this high accuracy is driven by VIS2, which represents higher-order visual areas, with less contribution from VIS1, representing low-order visual areas.

**Fig. 5.**
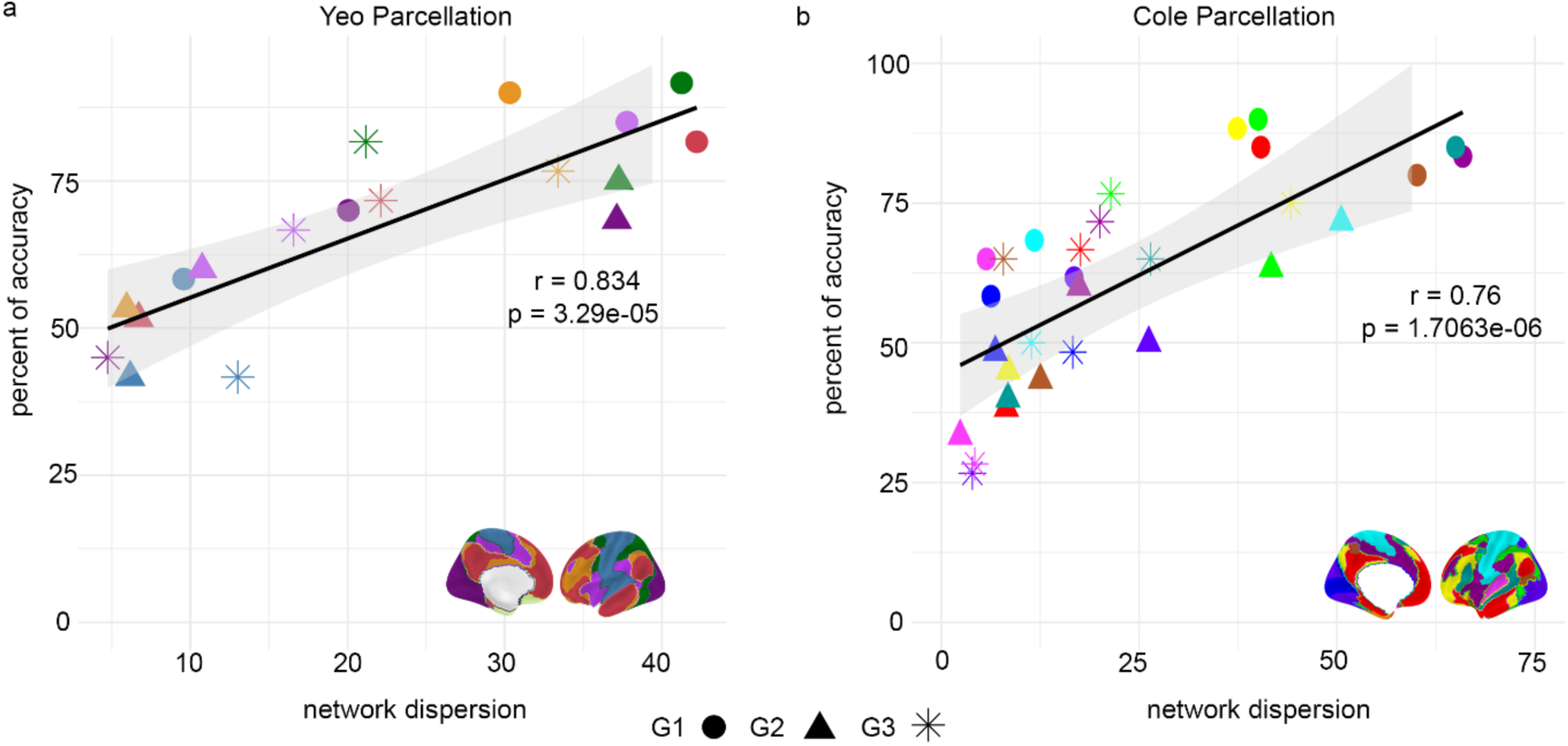
Identity accuracies in networks positively correlate with their dispersion along the gradients. Each point represents a different network, with accuracy rates on the y-axis and network dispersion on the x-axis. **(a)** shows results for the Yeo parcellation (Thomas Yeo et al., 2011), **(b)** shows results for the Cole parcellation (Ji et al., 2018). In both parcellations, higher network dispersion is associated with higher accuracy rates, indicating that a broader range of connectivity leads to a more predictive network of individually unique organization. Brain maps illustrate corresponding networks for each parcellation.

### Networks’ dispersion is associated with higher accuracy rates in identification analysis

To investigate how dispersion is related to accuracy rates in the identification analysis, we correlated accuracy rates in specific networks with their dispersion as measured by the variance of loading values across the three different connectivity gradients. The results reveal a significant positive correlation between these two variables. Specifically, as network dispersion increases, accuracy increases, indicating that a broader range of connectivity contributes to more stable network loading coefficient values over time. This relationship was consistent across both the Yeo (r = 0.834, p = 3.29e-05) and Cole (r = 0.76, p = 1.706241e-06) parcellation schemes while accounting for the number of vertices in each network by means of partial correlation.

## Discussion

Here we aimed to assess the stability of connectivity gradients along the first three connectivity gradients and to determine if they can be used for effective connectome fingerprinting over short and long-term time intervals. We additionally investigated how connectome fingerprint accuracies are influenced by the complexity of networks along the hierarchy and the dispersion of their gradient profiles.

We found that the first three connectivity gradients demonstrated high stability over time as measured by spatial correlation. Additionally, connectome fingerprinting based on correlation values demonstrated significantly high accuracy rates in all three connectivity gradients. Observed accuracies were slightly higher for the first and the third connectivity gradient than for the second connectivity gradient. Importantly, although a certain reduction in correlation values and accuracy rates was found over time, effective fingerprinting was still retained even when scans were separated by a one-month period, further strengthening the stability of connectivity gradients within individuals and the utilization of the approach for representing unique brain organization.

When using a-posteriori defined parts of the gradients as a basis for connectome fingerprinting (Ji et al., 2018; Thomas Yeo et al., 2011), we found that the complexity of networks was associated with accuracy rates such that heteromodal networks showed higher accuracies as compared with unimodal networks. This result was observed along the first and the third gradient, but not the second gradient. A closer look into both the dispersion of gradient coefficients of the different networks and the specific emphasized representation along the second connectivity gradient provided an explanation for this result. The *gradient dispersion* measured by the variance of values in specific networks showed a significant positive correlation with the fingerprinting accuracy rates. This relationship was evident taking all gradients into account and using two different parcellations. In terms of representation, the second connectivity gradient demonstrates the dissociation between the visual and the somatosensory networks. By using a more detailed parcellation, a difference emerged between high-order and low-order visual networks such that fingerprinting accuracies were higher in more complex visual areas.

Taken together with the results of the first and the third gradient this suggests that network complexity and importantly its dispersion along the gradients dictate fingerprinting performance.

Beyond the unique organization captured along the first three gradients, information contained in the gradients is also largely shared across participants. This is the case along the first and the second connectivity gradients showing high correlation values. In contrast, along the third connectivity gradient correlation values drop and are more moderate. When looking at the ratio of within to between subjects’ similarity, we found that similarity within subjects was always higher than similarity between subjects setting average ratio values above one. Interestingly, along the third gradient, within subjects’ similarity drops but to a lesser extent than between subjects’ similarity, significantly increasing the ratio as compared with the first and the second gradients. This suggests that the third gradient becomes more variable across individuals.

Our results replicate previous work investigating test-retest reliability of the first connectivity gradient in different datasets (Hong et al., 2020; Knodt et al., 2023). Hong et al., have examined how variations in preprocessing influences reliability for the first 100 gradients (Hong et al., 2020), while more recently, Knodt et al., have directly compared reliability based on the first connectivity gradient to that of edge-wise functional connectivity measures (Knodt et al., 2023). Both studies found good reliability for the first connectivity gradient (Hong et al., 2020; Knodt et al., 2023). Our study further extends previous work by investigating both stability in the first three gradients beyond the principle one and the associated connectome fingerprinting accuracy rates (Finn et al., 2015; Ramduny & Kelly, 2024) spanned over short and long-term intervals. Connectome fingerprinting provides an intuitive and easily interpretable measure of connectome stability over time. By effectively demonstrating an individual can be identified in a group, numeric measures of agreement are complemented by a straightforward quantification of repeated measures stability (Ramduny & Kelly, 2024). Our within subjects correlation values were slightly higher than those reported in Knodt et al., yet this can be explained by both the algorithm used for dimensionality reduction (PCA used here versus diffusion embedding used in Knodt et al.,) which has shown higher reliability values in previous work (Hong et al., 2020) and an a-priori spatial down-sampling which was not implemented in our study yet is often implemented in gradients analysis (Vos de Wael et al., 2020). Retaining all vertices may thus improve reliability and associated accuracy rates for individual connectome fingerprinting.

The slight differences found here in terms of higher accuracy rates reported for the first and the third connectivity gradients can be linked with previous work emphasizing these two gradients in detecting the spread of individual lesions within the functional connectome thereby suggesting stronger individual differences may be carried by these two gradients (Bayrak et al., 2019). Combined with the more variable ratio of within between reliability along the third gradient, this suggests combining the first and the third connectivity gradients may benefit analyses of individual behavioral predictions, however this is yet to be determined by future studies. Taken together, our findings strengthen the usage of the first three connectivity gradients as measures of individual brain organization stable over time. Overall, previous work establishing a link between connectivity gradients, individual behavior and symptomatology along mostly the first (Dong et al., 2023; Girn et al., 2022; Hong et al., 2019; Meng et al., 2021; Xia et al., 2022), but also secondary gradients (Bethlehem et al., 2020; Brown et al., 2022; Girn et al., 2022; Setton et al., 2022; Sydnor et al., 2021) further supports the functional relevance of the approach for prediction of individual cognitive behaviors.

Beyond demonstrating that connectivity gradients characterize individual features, information on the features that contribute to this uniqueness can be additionally drawn from our study. Different canonical networks are differently positioned along the continuum of the first three gradients. Additionally, the *dispersion of* network-specific gradient coefficients changes as well. Depending on the representation captured in a specific gradient, network dispersion varies, thereby enabling us to determine if it is associated with fingerprinting accuracy rates and how it relates to the complexity of the network. Previous work based on parcellation studies has established that specific networks are more predictive than others in terms of individual uniqueness. While there is some discrepancy, it is largely agreed that networks of higher complexity (i.e., heteromodal) and specifically frontal and default-mode networks are the most predictive of individual uniqueness (Blautzik et al., 2013; Marchitelli et al., 2017; Mejia et al., 2018; Noble et al., 2017, 2019; O’Connor et al., 2017; Somandepalli et al., 2015; Wisner et al., 2013; Zuo et al., 2014). Here, we replicate this result by showing that loading values in heteromodal networks yield higher accuracy rates as compared with unimodal networks along the first and the third connectivity gradients, but not the second. This result highlights the importance of network complexity, or its hierarchical role. However, network positioning and its dispersion changes along different gradients. This creates a situation where a given network with a well-known hierarchical role can change its *range* in different gradients thereby reflecting a different element of its representation. Indeed, along the second gradient the dissociation between unimodal and heteromodal in terms of accuracy seems to fall out. However, a closer look is showing that although the visual network increases its accuracy, this result is driven by high-order visual areas. Overall, we found that the dispersion of a given network is associated with accuracies irrespective of location along the different gradients. This result is in accordance with Knodt et al., where range has been shown to be associated with higher ICC values (Knodt et al., 2023), as well as theoretical and evolutionary accounts of the ‘untethering’ of high-order cognitive areas from sensory ones (Buckner & Krienen, 2013) and the distance between them as reflecting the flexible range that affords abstract cognitive functions (M. M. Mesulam, 1998). Taking our results together with previous studies emphasizes that the dispersion, reflecting the degree of connectedness of a given network (van den Heuvel & Sporns, 2013), is an important factor influencing uniqueness in addition to complexity. Our result can explain some discrepancies in parcellation based studies showing visual networks are highly reliable within individuals (Blautzik et al., 2013; B. Chen et al., 2015; Noble et al., 2019; Shirer et al., 2015), and crucially stresses that the degree of network’s integration is an important factor, even if not always independent, accompanying network complexity along the hierarchy in the prediction of individual brain signatures. It remains to be determined in future studies what is the broader utility of dispersion as a biomarker of individual behaviors.

Investigating the spatial similarity between different subjects complements information on unique features of brain organization with those that are mutual. The high correlation values between subjects especially along the first and the second gradient suggests the representation captured in these gradients is mostly shared across people, promoting the adequacy of using the approach for group comparisons. This result is in line with studies demonstrating gradients are resilient to state-dependent changes following sleep deprivation (Cross et al., 2021) along with studies in monkeys demonstrating a similar organization (Buckner & Margulies, 2019). Additionally, group studies that show differences in connectivity gradients, even in pathological versus healthy population (Dong et al., 2023; Hong et al., 2019; Meng et al., 2021; Xia et al., 2022) are expressing these differences in ‘contraction’ or ‘expansion’ of the gradients and not in dismantling of the order retained in the representation. Additional more methodological studies demonstrate the organization is preserved to a large extent even when different algorithms (Hong et al., 2020) or different thresholding strategies are used (Nenning et al., 2023). More generally, the representation captured along the first connectivity gradient is spanning two, largely antagonistic systems at its extremes. On the one end an intrinsic system dealing with our internal world and more abstract cognitive functions, and on the other end an extrinsic system dealing with our sensory world (Buckner et al., 2008; Buckner & Krienen, 2013; Golland et al., 2007; Raichle & Snyder, 2007; Smallwood et al., 2021). It is interesting to note that beyond the overt cognitive dissociation of these two systems, they tend to repeatedly emerge as separate entities in data driven analyses (Pines et al., 2022) thereby further supporting this representation as a fundamental organizing principle.

Our results should be interpreted considering several limitations. First, connectivity gradient analysis usually involves alignment of individual gradients to promote adequate comparison and mutual order of the gradients (Langs et al., 2015; Vos de Wael et al., 2020). As the approach is data-driven and the order of gradients is a byproduct of the explained variance, some variability exists in the order of gradients between individuals. Here, we did not examine how variable this factor is. Although alignment is a very common and acceptable practice in the field, future work should examine more systematically if individual order of gradients is in any way informative for individual fingerprinting. Second, our analysis was restricted to the first three connectivity gradients. This decision was driven by both the cumulative explained variance saturating quite rapidly after the first three gradients, in addition to the interpretability challenge associated with increased number of gradients, and the majority of brain-behavior studies using the approach focusing on the first, or the first several gradients (Bethlehem et al., 2020; Brown et al., 2022; Dong et al., 2023; Girn et al., 2022; Hong et al., 2019; Meng et al., 2021; Setton et al., 2022; Sydnor et al., 2021; Xia et al., 2022). However, recent work showing gradient-behavior prediction can improve using more than several gradients (Kong et al., 2023) suggest that more gradients may be relevant for predicting individual behavior. Lastly, we defined short- and long-term intervals as separated by three days and 30 days, respectively. This with the logic that individual signatures should be stable over long time periods, and the expected decay in stability over time. Future work should investigate the stability of gradients over longer time scales using repeated scanning spanned over longer time periods to support this claim (Laumann et al., 2015; Poldrack et al., 2015).

## Conclusions

We have shown that the first three connectivity gradients are stable over time within individuals both on short and long-term intervals. Stability was accompanied by high accuracy rates demonstrating connectivity gradients can be effectively used for connectome fingerprinting purposes. When examining which features contribute to the individual uniqueness, we found that the dispersion of networks, along with their complexity in terms of hierarchical role are related to accuracy such that the higher the complexity is, and importantly the higher the dispersion is, the better accuracy rates are. This result was evident even when taking differences in network sizes into account, in two different parcellations, and importantly along the three different gradients where networks are located in different places on the continuum and can show differences in dispersion. Beyond the uniqueness, the first and the second connectivity gradients also showed high similarity between individuals demonstrating the organization captured along these two gradients is robust across participants and largely shared. Our study extends previous work on the shared versus unique organizational features and offers insights into the importance of network dispersion, i.e., the degree to which a given network shares connectivity with other networks, to the individual uniqueness it carries.

## Acknowledgements

The authors would like to kindly thank Dr. Yating Lv and Dr. Bella Vakulenko-Lagun for their advice on statistics and preprocessing. This work was supported by the Max-Planck Partner Group funding scheme; by the National Institute for Psychobiology in Israel Founded by the Charles E. Smith Family (Grant Number 25-20-21); and by the Alon Scholarship for the Integration of Outstanding Faculty (Israeli Council for Higher Education). Grants were obtained by Dr. Ovadia-Caro.

