## Supplemental materials for "Individual uniqueness of connectivity gradients is driven by the complexity of the embedded networks and their dispersion"

**Supplementary Materials**

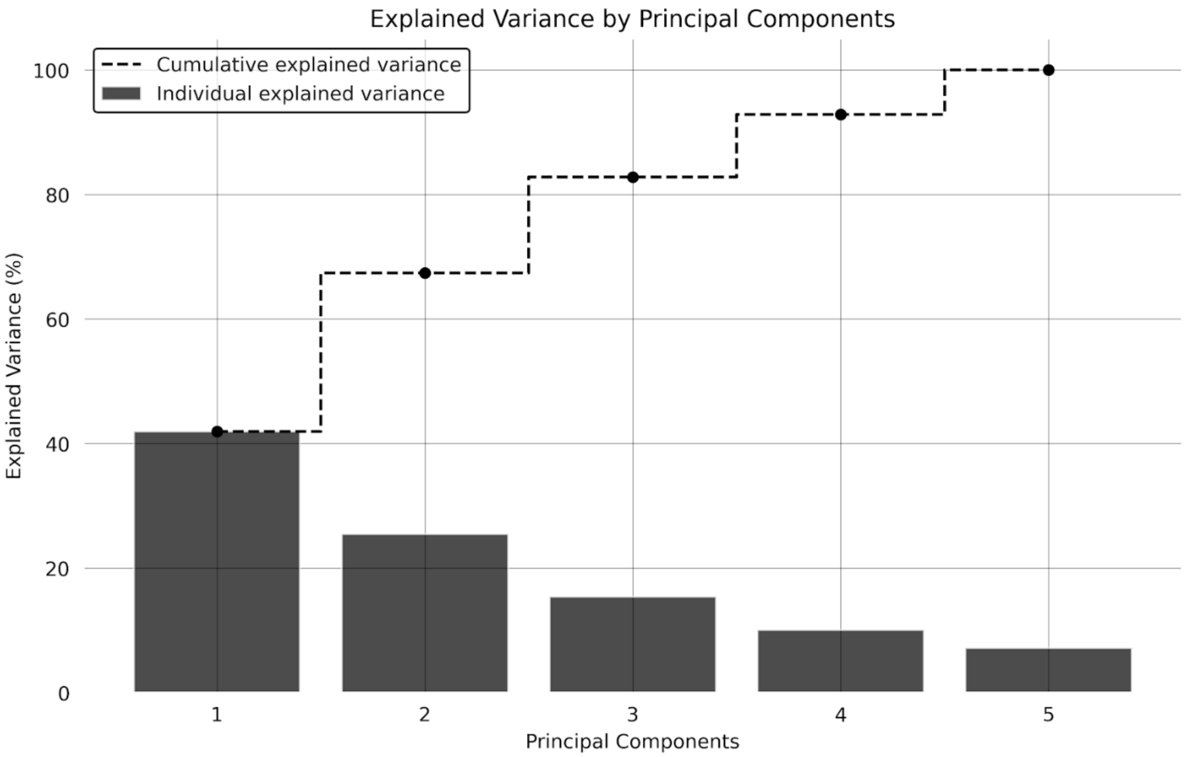


***Fig. S1 Individual and cumulative explained variance by connectivity gradients***. Percent explained variance for each component out of the 5 components decomposed in our analysis. The dashed line indicates the cumulative explained variance. The principal component accounts for the largest proportion of variance, at 42%. Notably, the cumulative explained variance approaches 85% by the third component.
